# Cannabidiol reverses microglia activation and loss of parvalbumin interneurons and perineuronal nets in a mouse model of schizophrenia

**DOI:** 10.1101/2024.10.21.619352

**Authors:** Naielly Rodrigues da Silva, Davide Gobbo, Felipe V. Gomes, Anja Scheller, Frank Kirchhoff, Elaine Del Bel, Francisco Silveira Guimarães

**Affiliations:** Department of Pharmacology, Ribeirao Preto Medical School, University of Sao Paulo, Ribeirao Preto, Brazil; Molecular Physiology, Center for Integrative Physiology and Molecular Medicine (CIPMM), University of Saarland, Homburg, Germany; Department of Basic and Oral Biology, Faculty of Odontology of Ribeirão Preto, University of São Paulo, Ribeirão Preto, Brazil

**Author notes:** Corresponding Author: Naielly Rodrigues da Silva, Department of Pharmacology, Ribeirão Preto Medical School, University of Sao Paulo, 3900 Bandeirantes Avenue, Ribeirao Preto, SP, 14049-900, Brazil.

**Keywords:** cannabidiol, antipsychotic, parvalbumin interneurons, microglia, NMDA hypofunction

## Abstract

Cannabidiol (CBD) has shown potential for treating schizophrenia (SCZ) by targeting its positive, negative, and cognitive symptoms. In this study, we investigated if CBD could reverse the memory impairment observed after chronic administration of the NMDA receptor antagonist. MK-801 treatment (0.5 mg/kg i.p., twice a day, for 14 days) resulted in short- and long-term memory deficits and decreased relative power of γ oscillation in freely moving animals. CBD administration (60 mg/kg i.p. daily for seven days after the MK-801 treatment period) reversed these changes. The positive cognitive effects of CBD were prevented by a 5-HT1A, but not a CB2, receptor antagonist. On the cellular level, CBD reversed MK-801-induced reduced number of parvalbumin-positive neurons and their associated perineuronal nets in the prelimbic medial prefrontal cortex (mPFC) and ventral hippocampus (vHip). This neuroprotective effect was mediated by 5-HT1A and CB2 receptors in the vHip but was independent of these receptors in the mPFC. Additionally, CBD reversed MK-801-induced microglial activation in both mPFC and vHip, again through 5-HT1A and CB2 receptors. These findings suggest that CBD modulates multiple pathways affected in SCZ-like conditions, offering a promising therapeutic avenue for SCZ treatment.

**Chemical compounds studied in this article:** CBD (PubChem CID: 644019), MK-801 (PubChem CID: 180081), AM630 (PubChem CID: 4302963), WAY100635 (PubChem CID: 11957721).

## Introduction

Schizophrenia (SCZ) is a complex, chronic, and disabling disorder with an elusive underlying neurobiological mechanism [1]. Symptoms of this disorder typically include positive (e.g., delusions and hallucinations), negative (e.g., anhedonia and social isolation), and cognitive symptoms (e.g., impaired verbal learning, executive function, and working memory) [2-4].

A hypofunction of glutamate N-methyl-d-aspartate receptors (NMDAR) is proposed to underlie a wide range of SCZ symptoms [5-6]. This idea derives from clinical and preclinical studies showing that acute and chronic administration of the NMDAR antagonists induces behavioral changes similar to the positive, negative, and cognitive symptoms of SCZ [7]. Interestingly, in animal models, the behavioral deficits induced by repeated administration of NMDAR antagonists can last up to 6 weeks [8]. In addition to behavioral abnormalities, NMDAR antagonists induce neurochemistry and neuroanatomical changes related to SCZ [9-11].

It has been suggested that the impact of NMDAR antagonists is more significant on fast-spiking parvalbumin-positive (PV^+^) GABAergic interneurons [6]. PV^+^ interneurons play a critical role in controlling the rhythmic activity of pyramidal neurons, which is essential for generating brain γ oscillations in sensory and cognitive processes that are impaired in SCZ [12-14]. A significant reduction in the number of PV^+^ interneurons has been consistently found in the prefrontal cortex (PFC) and hippocampus of post-mortem brains of SCZ patients [15-17] and in several rodent SCZ models [18]. PV^+^ interneuron hypofunction is proposed as a critical etiologic factor of SCZ [19]. The hypofunction of NMDAR-mediated signaling could be involved in the observed PV^+^ neuronal dysfunction in SCZ [6, 20-22].

Most of the PV^+^ interneurons in the brain are surrounded by the perineuronal nets (PNNs), a specialized extracellular matrix that aggregates around the neuronal soma and is directly involved in the regulation of PV^+^ interneurons synaptic plasticity [23], as well as in their protection against oxidative and metabolic damage [24, 25]. PNNs mature gradually, in an experience-dependent manner, during the maturation of neural circuits in early adulthood, coinciding with the age of SCZ onset [26]. Decreases in PNNs, including those surrounding PV^+^ interneurons, have also been observed in SCZ patients [27] and animal models of the disease [28]. However, the mechanisms underlying PNN deficits in SCZ, which may include the functional capacity of microglial cells to remodel the brain extracellular matrix [23, 29, 30], are still unclear.

Current therapy for SCZ is based on antipsychotic drugs that often cause limiting side effects and fail to alleviate the negative and cognitive symptoms, which are strongly predictive of poor functioning and long-term outcomes [30-33]. Clinical and preclinical studies point to the possible antipsychotic properties of cannabidiol (CBD), the major non-psychotomimetic compound present in the *Cannabis sativa* [34, 35]. Our group has reported that repeated CBD treatment prevented and reversed the social and cognitive impairments in a mouse model of SCZ based on chronic treatment with the NMDAR antagonist MK-801 [9, 36, 37]. Also, CBD, similar to the second-generation antipsychotic clozapine, prevented the decrease in the number of PV^+^ interneurons in the PFC [36].

However, CBD’s impact on the loss of PV^+^ interneurons and PNNs in SCZ models is still poorly explored. Also, even if in our previous work CBD (when administered concomitantly with MK-801) prevented the microglia activation caused by repeated administration of an NMDAR antagonist, it is unknown if it could reverse these findings. Concerning the potential mechanisms of CBD antipsychotic effects, the potentiation of endocannabinoid signaling by inhibiting the fatty acid amide hydrolase (FAAH), which hydrolyzes anandamide (AEA), has been proposed [38, 39]. In line with this hypothesis, the clinical improvement observed in SCZ patients after CBD treatment correlated with increases in AEA plasma levels [40]. Also, CBD decreases MK-801-induced microglial activation [36]. This latter effect could be mediated by CB2 receptors, whose activation is known to decrease microglia-mediated neurotoxicity and pro-inflammatory cytokine levels [41, 42]. Another mechanism that could be associated with CBD antipsychotic action is the facilitation of serotonin-1A (5-HT1A)-mediated neurotransmission [37]. Since 5-HT1A receptors control neuronal firing in the PFC [43-46], possibly via PV^+^ interneurons [47], it is of particular interest to investigate the contribution of 5-HT1A receptors in the antipsychotic effect of CBD [36, 48].

Therefore, this work investigated the potential involvement of PV^+^ interneurons, PNNs, and microglia activation on CBD antipsychotic effects in a SCZ model based on chronic NMDAR antagonism. We also investigated if these effects depend on CB2 and/or 5-HT1A receptors.

## Materials and Methods

### Animals

6-week-old male C57BL/6J mice were used in the experiments. They were housed in groups of five per cage (41 x 33 x 17 cm) in a temperature-controlled room (24 ± 1°C) under standard laboratory conditions with free access to food and water and a 12 h light/dark cycle (lights on at 6:00 AM). Procedures were conducted in conformity with the Brazilian Society of Neuroscience and Behavior guidelines for the care and use of laboratory animals, which follow international laws and politics. The Animal Ethics Committee of the institution approved the housing conditions and experimental procedures (process number 145/2015). Adequate care has been taken to minimize pain, discomfort, or stress to the animals, and every effort has been made to use the minimum number of animals necessary to obtain reliable scientific data.

### Drugs

The following drugs were used: CBD (BSPG, UK) diluted in 1% DMSO + 2% Tween 80 in saline, MK-801 (Sigma-Aldrich, USA) diluted in saline, AM630 (CB2 receptor antagonist, Sigma-Aldrich, USA) diluted in 1% DMSO + 2% Tween-80 in saline, and WAY100635 (5-HT1A receptor antagonist, Tocris, USA) diluted in saline. The drugs were injected intraperitoneally (i.p.) in a 10 mL/kg volume.

### Pharmacological treatment

To assess the effect of CBD and the involvement of 5-HT1A and CB2 receptors in CBD-mediated effects on MK-801-induced cognitive impairment, 6-week-old male C57BL/6J mice were treated with MK-801 (0.5 mg/kg, 14 days, 2x/days) or vehicle twice a day for 14 days. One day after the end of treatment with MK-801, the mice were injected i.p. with the 5-HT1A receptor antagonist WAY100635 (0.1 mg/kg, seven days, 1x/day), the CB2 receptor antagonist AM630 (0.1 mg/kg, seven days, 1x/day) or vehicle. Each injection was followed, 10 min later, by CBD (30 mg/kg, i.p., seven days, 1x/day) or vehicle. One day after the end of treatment, the novel object recognition (NOR) test was performed. To assess the effect of CBD on MK-801-induced alterations in the electroencephalogram (EEG), 6-week-old male C57BL/6J mice were treated with MK-801 (0.5 mg/kg, 14 days, 2x/days) or vehicle twice a day for 14 days. On day 7 of the treatment, mice underwent surgery for telemetric EEG recording. One day after the end of treatment with MK-801, the recording started, and the mice were treated with CBD (30 mg/kg, seven days, 1x/day) or vehicle. The mice were perfused for tissue collection and immunohistochemistry at the end of the procedure. The treatment regimen and doses used were based on previous studies from our group [37].

### Novel object recognition (NOR) test

The NOR test was carried out in a circular arena (40 cm high and 40 cm in diameter). Each animal was submitted to a habituation session in the arena (15 min) and exposed the following day to two identical objects in the same arena (acquisition, 10 min). After 60 min, one of the objects was replaced by a new one, and the animals were placed in the arena containing the new and familiar objects (NOR-ST, retention short-term memory [NOR-ST], d2, 5 min). The following day, a new object replaced the first one (retention long-term memory [NOR-LT], 5 min). The discrimination index (DI) was used to evaluate the relative exploration time of the two identical objects during the acquisition phase and the new and familiar objects during the retention phases. It was calculated as DI = (T1 *or* N – T2 *or* F)/(T1 *or* N + T2 *or* F), where T1 and T2 are the time of exploration of the two identical objects in the acquisition and TN and TF are the new and familiar objects’ exploration time, respectively.

### Telemetric EEG implantation and recording

Mice were implanted with telemetric EEG transmitters (DSI PhysioTel ETA-F10, Harvard Biosciences, USA) to allow telemetric EEG recording in their home cage. The animals were placed in a stereotaxic frame (Robot stereotaxic, Neurostar, Germany) to implant depth electrodes at 3.4 mm posterior to bregma and bilaterally 1.6 mm from the sagittal suture. After postsurgical care and recovery, cages were placed on individual radio receiving plates (DSI PhysioTel RPC-1, Data Sciences International, USA), which record EEG signals and send them, together with the video recording (MediaRecorder Software, Noldus Information Technology, Netherlands) to an input exchange matrix (DSI PhysioTel Matrix 2.0 (MX2), Ponemah software, DSI, Data Sciences International, USA). EEG recording started 2 hours before the first CBD injection (baseline) and throughout the treatment with CBD. EEG traces were analyzed with the Neuroscore software (Version 3.3.1., Data Sciences International, USA). Data are displayed as average power spectrum in the specific band range within the period evaluated.

### Immunohistochemistry

The mice were anesthetized with a lethal dose of urethane (25 %, 5 mL/kg) and submitted to transcardiac perfusion with a volume of 30 mL/animal of a phosphate saline solution (PBS, 0.01 M, pH 7.4) followed by 25 mL/animal of paraformaldehyde fixative (4 % PFA) in phosphate buffer (0.1 M PB, pH 7.4). The brains were extracted and kept soaked in the fixative (4% PFA) for 24 h, cryoprotected with 30 % sucrose in 0.01 M PBS for 72 h at 4°C, frozen with isopentane at -40°C in dry ice, and kept stored until processing at -80°C. Adjacent 30 µm-thick coronal slices were obtained using a cryostat (CM-1900, Leica, Germany) at -24°C and stored at -20°C in an antifreeze solution.

For immunohistochemistry, the sections were washed in 0.01 M PBS and 0.1 % Triton-X-100 (Buffer A, 3x, 5 min each) and incubated with citrate buffer (pH 6) for 30 min at 60°C. After washing (Buffer A, 3x, 5 min each), the sections were blocked for two h in Buffer A with 5 % bovine albumin serum (BSA) and incubated overnight at 4°C in slow agitation with primary antibody and/or biotinylated Wisteria Floribunda lectin (WFA, 1:1000, Vector Labs, USA, Ref. B-1355-2). The following primary antibodies were used: rabbit anti-parvalbumin (1:1000, Cell Signaling, USA; Ref. 55598) and rabbit anti-Iba1 (1:500, Invitrogen, USA Ref. MA5-41239). After washing (Buffer A, 3x, 5 min each), the sections were incubated for 2 hours with Alexa Fluor 649 anti-rabbit (1:500, Invitrogen, USA). Afterward, the sections were washed (Buffer A, 3x, 5 min each), mounted on gelatinized glasses protected from light, and dried for 8 hours. Slides were mounted with a mounting medium containing DAPI (Abcam, USA; Ref. ab104139) and kept at 4°C until analysis.

### Immunohistochemistry analysis

Higher magnification acquisitions were obtained using a confocal microscope (Leica SP5) (PV/WFA, 20x magnification; Iba1, 40x magnification). Acquisitions were recorded from the prelimbic and infralimbic regions of the medial PFC (mPFC) and ventral CA1 and ventral subiculum (vSub) of the hippocampus. Areas were identified according to the Paxinos and Franklin Mouse Brain Atlas [71]. Six tissue sections were collected per area of interest and animal. Z-stack maximum projections were obtained and analyzed using ImageJ® software (NIH, USA) for fluorescence intensity measurements. Microglia number was assessed using the ImageJ Cell Counter plugin.

### Statistical analysis

The data are presented as mean ± standard error of the mean (SEM) and were analyzed by Student’s t-test or one-way analysis of variance (ANOVA). Post-hoc comparisons were made using the Dunnett or Student-Newman-Keuls (S-N-K) test. A significance level of 95 % (p<0.05) was used, and the analysis was carried out using IBM SPSS software (version 21.0).

## Results

### CBD counteracts SCZ-like cognitive disruption through 5-HT1A receptors

To model SCZ-like cognitive symptoms, we treated mice with the NMDAR antagonist MK-801 for 14 days (d0-d13). We tested for short- and long-term memory impairments one week after the end of the MK-801 treatment (d21-23) (Figure 1A). No difference was observed in the exploration of the two identical objects during the acquisition phase of the NOR test for any group tested (Figure 1B and Supplementary Figures 1A-B). In the short-memory retention session (NOR-ST), the time spent exploring the novel object compared to the familiar object was unaltered in mice treated with MK-801, indicating a failure to discriminate between familiar and new objects. This impairment was counteracted by CBD since MK-801-treated mice administered with CBD (d14-d20) spent more time exploring the new object, which was associated with a positive discrimination index. CBD effects were prevented by the 5-HT1A receptor antagonist WAY100635 but not the CB2 receptor antagonist AM630 (Figure 1C and Supplementary Figure 1C). Similar results were obtained the following day in the long-term memory retention session (NOR-LT, Figure 1D and Supplementary Figure 1D).

**Figure 1.**
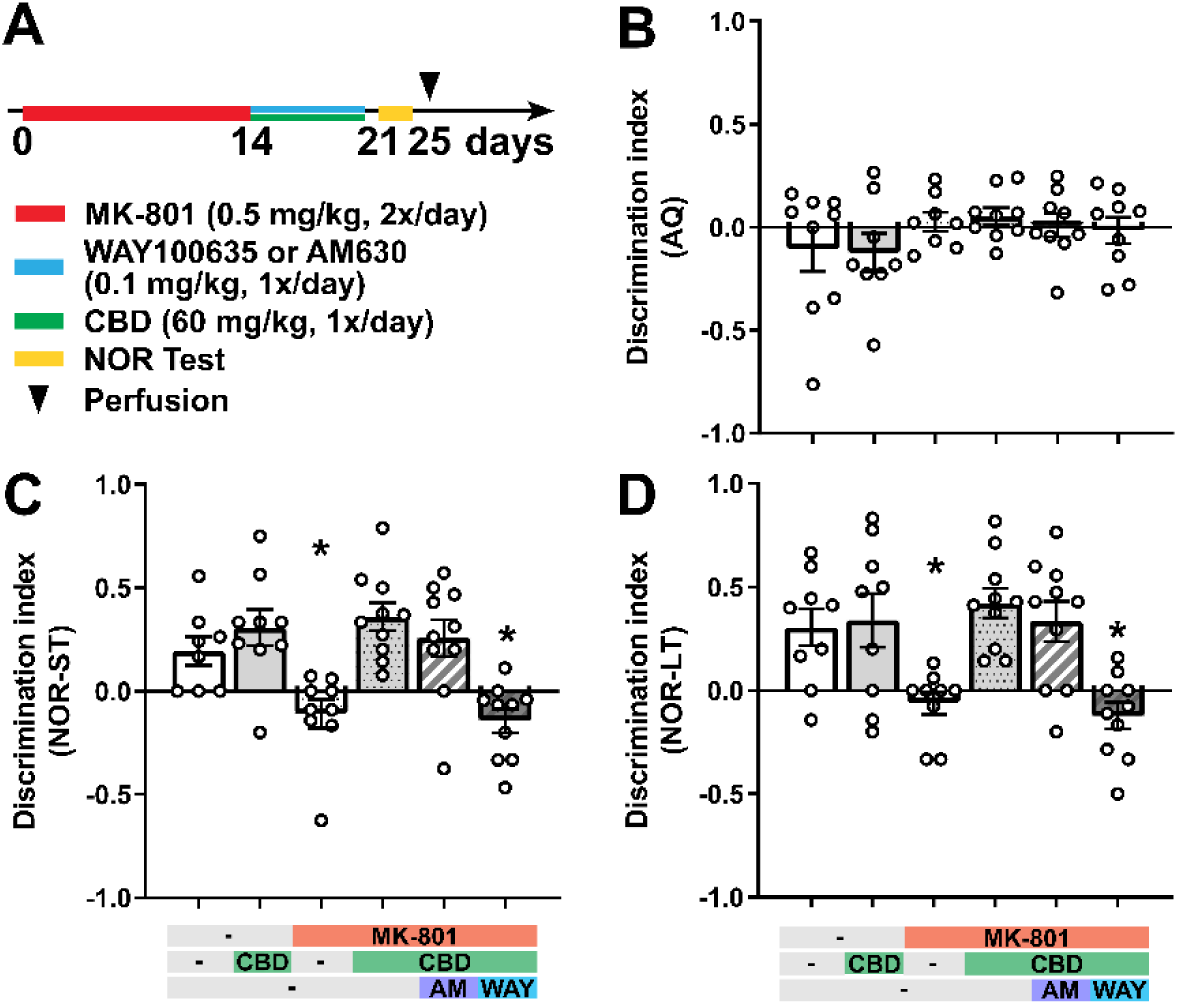
CBD treatment reduces schizophrenia-like cognitive symptoms through 5-HT1A receptors. **(A)** Experimental protocol for pharmacological treatment with the NMDA receptor antagonist MK-801 (d0-d13), WAY100635, or AM630 and CBD (d14-d20) followed by novel-object-recognition test (habituation, d21; acquisition and retention, d22-23). **(C-D)** Discrimination index [calculated as (T_N_ - T_F_)/(T_N_ + T_F_), where T_N_ and T_F_ are the exploration time of the new and the familiar object, respectively] during the short-term memory retention phase (NOR-ST, **C**) and long-term memory retention phase (NOR-LT, **D**). During the acquisition phase (AQ, **B**), the results are presented as a discrimination index between the two identical objects. Data are represented as mean ± SEM derived from 8-9 mice. Each group was compared to the control group (*, p < 0.05). Details on statistical analysis can be found in Supplementary Table 1.

### CBD prevents changes in γ oscillation associated with working memory impairment

The impairment of cognitive processes associated with SCZ is linked to alteration in the function of cortical PV^+^ interneurons, which play a critical role in controlling the rhythmic activity of pyramidal neurons and brain γ oscillations [12-14]. To assess MK-801-induced spectral alterations in the γ range (25-80 Hz), we performed telemetric EEG recording on freely moving animals starting chronically injected with MK-801. The brain electrical signal was collected between the two brain hemispheres through deep electrodes inserted in the cortical layers of the brain occipital lobes. We detected a marked decrease (almost 20 %) in the relative power of γ oscillation. Moreover, the treatment with CBD (administered throughout the recording once per day) reverted the reduction in the relative power of the γ band (Figure 2).

**Figure 2.**
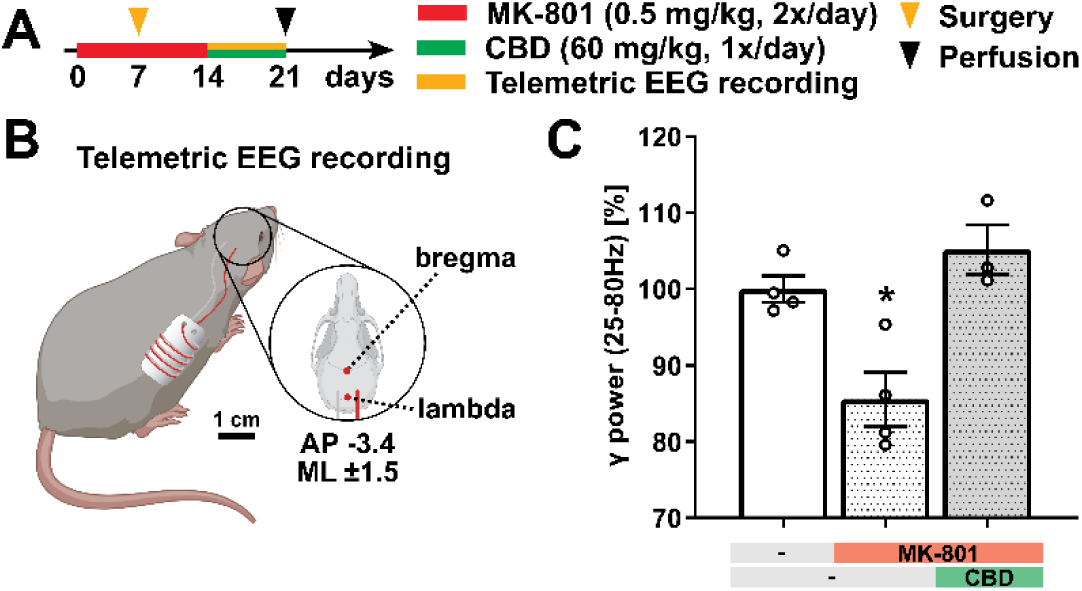
CBD treatment reverts MK-801-induced reduction of γ oscillation contribution. **(A)** Experimental protocol for pharmacological treatment with MK-801 and CBD and telemetric electroencephalographical recording (EEG) and perfusion. **(B)** Schematic representation of the ETA F-10 transmitter implanted to perform telemetric EEG recording. AP, anteroposterior axis; ML, mediolateral axis [Created with BioRender.com]. **(C)** The relative power of γ oscillations [25 - 80 Hz range] normalized on vehicle-treated mice calculated as average power within the period evaluated. Data are represented as mean ± SEM of 3-4 mice. Each group was compared to the control group (*, p < 0.05). Details on statistical analysis can be found in Supplementary Table 1.

### CBD protects against MK-801-induced loss of PV^+^ cells and PNNs in the prelimbic mPFC

We assessed the effect of MK-801 and CBD on PV^+^ cells and the integrity of WFA-stained PNNs in two different mPFC subregions, the prelimbic and infralimbic cortices (Figures 3A-C). In the prelimbic mPFC, MK-801 reduced the densities of PV^+^ cells and PV^+^ cells associated with PNNs (Figure 3F) and WFA density on PV^+^ cells (Figure 3H). Although CBD, by itself, also reduced the density of PV^+^ cells, it reverted the impairments caused by MK-801. This effect in the densities of PV^+^ cells and PV^+^ cells associated with PNNs was not blocked by AM630 and WAY100635 (Figures 3D and 3F). Regarding the infralimbic mPFC, no difference was observed in these variables (Figures 3E, 3G, and 3I). Also, no change was found in the total density of WFA^+^ cells or the total WFA density in the prelimbic and infralimbic mPFC (Supplementary Figure 2).

**Figure 3.**
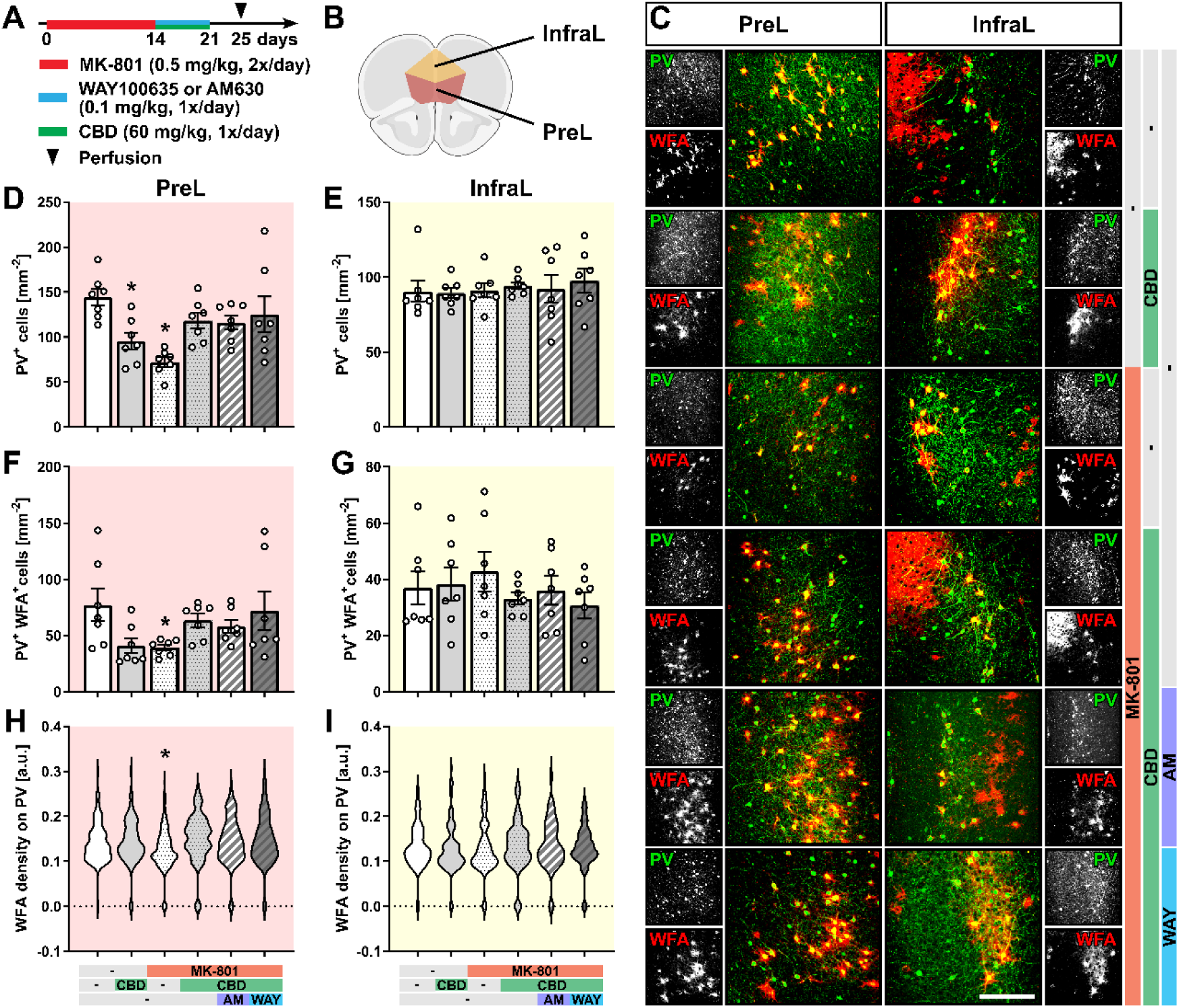
CBD protects against MK-801-induced decrease in the expression of PV^+^ neurons and PNNs in the prelimbic but not in the infralimbic region of the mPFC. **(A)** Experimental protocol for pharmacological treatment with MK-801, WAY100635, or AM630 and CBD followed by perfusion and immunohistochemical analysis. **(B)** Schematic representation of the prelimbic (PreL) and infralimbic regions (InfraL) of the mPFC used for analysis [Created with BioRender.com]. **(C)** A representative view of parvalbumin (PV, green) and PNNs was detected using WFA (red). Scale bar, 200 µm. **(D-E)** Cell density of PV^+^ cells in the PreL **(D)** and InfraL **(E)** regions. **(F-G)** Cell density of PV^+^/WFA^+^ cells in the PreL **(F)** and InfraL **(G)** regions. **(H-I)** WFA density (calculated as fluorescence intensity) on PV^+^ cells in the PreL **(H)** and InfraL **(I)** regions. Data are represented as mean ± SEM **(D-G)** or violin plot **(H-I)** and derived from N=7 mice **(D-G)** and 74-180 cells **(H-I)**. Each group was compared to the control group (*, p < 0.05). Details on statistical analysis can be found in Supplementary Table 1.

Taken together, these data suggest that MK-801-induced reduction in PV^+^ cell density and associated PNNs is restricted, in the mPFC, to the prelimbic cortex. CBD reverses this effect through mechanisms not involving 5-HT1A or CB2 receptors.

### MK-801-induced decrease of PV^+^ cells and PNNs in the vHip is reversed by CBD through 5-H1A and CB2 receptors

To evaluate the effect of MK-801 and CBD on PV^+^ cells and the integrity of WFA-stained PNNs in the vHip, we analyzed the ventral CA1 region (vCA1) as well as the ventral subiculum (vSub) (Figures 4A-C). Similar to the prelimbic mPFC, MK-801 reduced the densities of PV^+^ cells (Figures 4D and 4E) and PV^+^ cells associated with PNNs (Figures 4F and 4G), and WFA density on PV^+^ cells (Figures 4H and 4I) in these two areas. CBD prevented this effect. Contrary to the prelimbic region, CBD was blocked by AM630 and WAY100635 (Figures 4D-I). Like the mPFC region, the total density of WFA^+^ cells or the total WFA density in the vHip was substantially unaltered among the groups, although the total density of WFA^+^ cells reduced in the vCA1 of MK-801-treated mice (Supplementary Figure 3).

**Figure 4.**
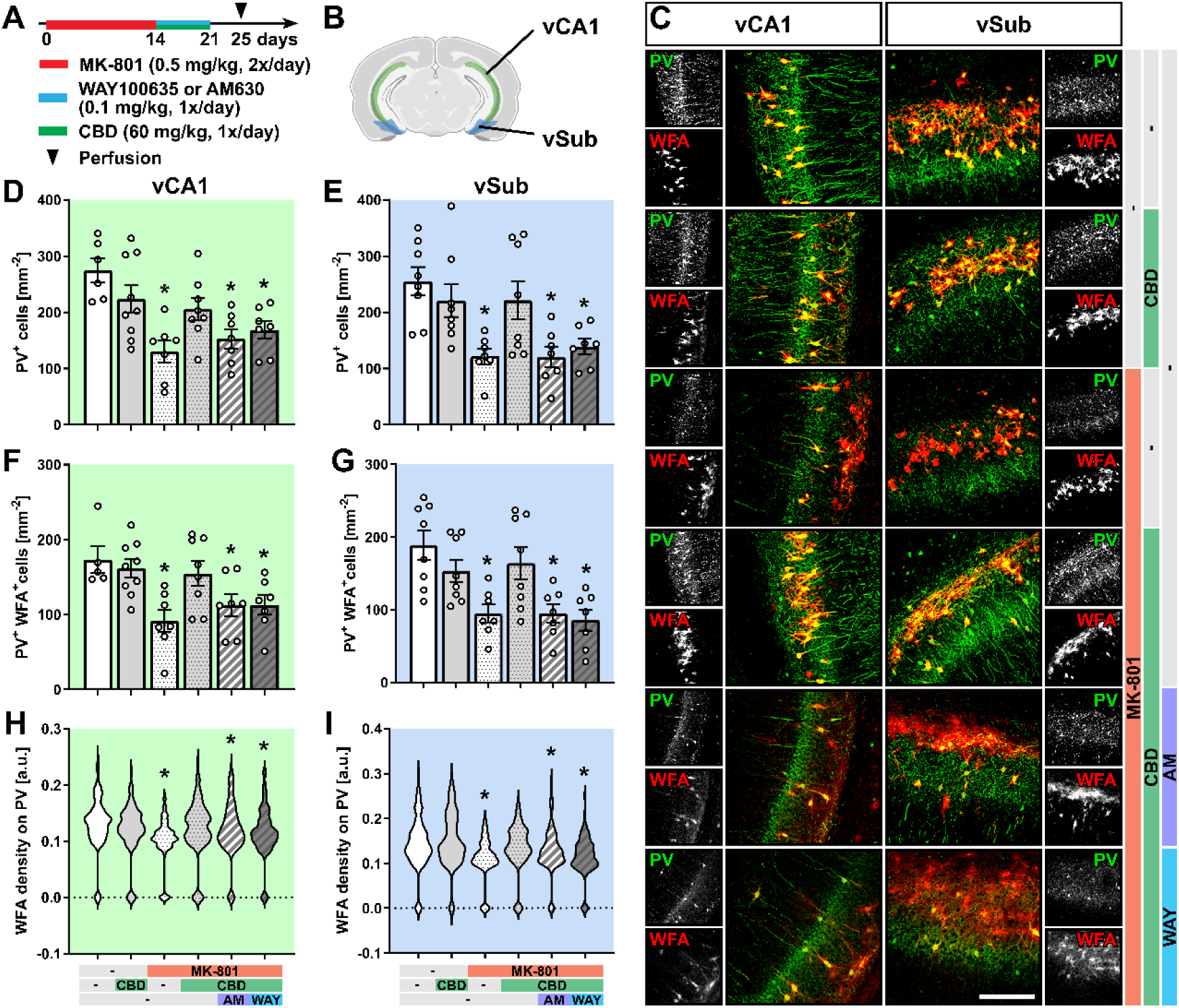
CBD protects against MK-801-induced loss of PV^+^ neurons and PNNs in the hippocampus through 5-H1A and CB2 receptors. **(A)** Experimental protocol for pharmacological treatment with MK-801, WAY100635, or AM630 and CBD followed by perfusion and immunohistochemical analysis. **(B)** Schematic representation of the ventral CA1 (vCA1) and the ventral subiculum (vSub) of the hippocampus used for analysis [Created with BioRender.com]. **(C)** A representative view of parvalbumin (PV, green) and PNNs was detected using WFA (red). Scale bar, 200 µm. **(D-E)** Cell density of PV^+^ cells in the vCA1 **(D)** and vSub **(E)** regions. **(F-G)** Cell density of PV^+^/WFA^+^ cells in the vCA1 **(F)** and vSub **(G)** regions. **(H-I)** WFA density (calculated as fluorescence intensity) on PV^+^ cells in the vCA1 **(H)** and vSub **(I)** regions. Data are represented as mean ± SEM **(D-G)** or violin plot **(H-I)** and derived from N=6-9 mice **(D-G)** and 90-199 cells **(H-I)**. Each group was compared to the control group (*, p < 0.05). Details on statistical analysis can be found in Supplementary Table 1.

These data suggest that MK-801 reduces PV^+^ cell density and associated PNNs in the vHip and that CBD reverses this effect by activating 5-HT1A and CB2 receptors.

### CBD reverses MK-801-induced microglia activation through 5-H1A and CB2 receptors in the mPFC and vHip

Next, we evaluated MK-801-induced microglial activation using the morphological classification of Iba1^+^ cells into five classes representing resting microglia (type I-II) and reactive microglia (type III-V) (Supplementary Figures 5A-B). In the prelimbic mPFC, MK-801 increased the percentage of reactive microglia. This result reflected a decreased proportion of type II resting microglia and an increased proportion of type III and IV reactive microglia. CBD treatment reverted the relative increase of reactive microglia, whereas the concomitant administration of AM630 or WAY100635 blocked the protective effect of CBD on MK-801-induced microglia activation (Figures 5A-D). Similar results were obtained in the infralimbic cortex, where MK-801 reduced the proportion of type II microglia and increased that of type III microglia. AM630 or WAY100635 blocked this effect (Figure 5E).

**Figure 5.**
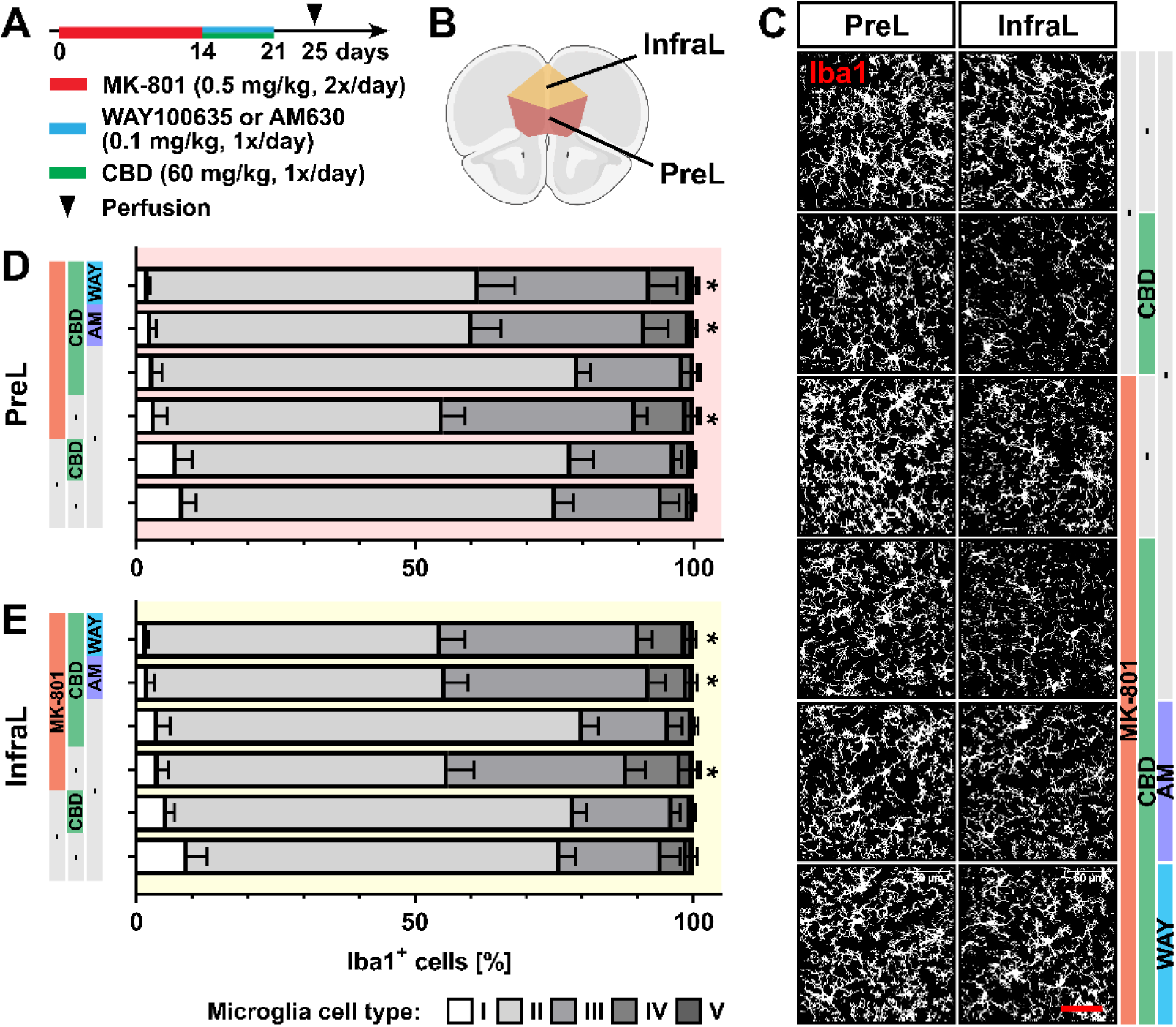
CBD reversed MK-801-induced microglia activation through 5-H1A and CB2 receptors in the mPFC. **(A)** Experimental protocol for pharmacological treatment with MK-801, WAY100635, or AM630 and CBD followed by perfusion and immunohistochemical analysis. **(B)** Schematic representation of the prelimbic (PreL) and infralimbic regions (InfraL) of the mPFC used for analysis [Created with BioRender.com]. **(C)** Representative view of Iba1^+^ microglia. Scale bar, 50 µm. **(D-E)** Relative number of microglia divided into five classes (resting microglia, type I-II; reactive microglia, type III-V) in the PreL **(D)** and InfraL **(E)** regions. Data are represented as mean ± SEM and derived from N=7-9 mice. Each group was compared to the control group (*, p < 0.05). Details on statistical analysis can be found in Supplementary Table 1.

Next, we evaluated microglia activation in the vHip (Figures 6A-C). Like the mPFC, MK-801 treatment increased the proportion of activated microglia in the vCA1 and vSub regions of the hippocampus, reducing type II resting microglia and increasing type III-IV reactive microglia. CBD blocked microglia activation induced by MK-801, and this effect was reverted by AM630 and WAY100635 (Figures 6D and 6E).

**Figure 6.**
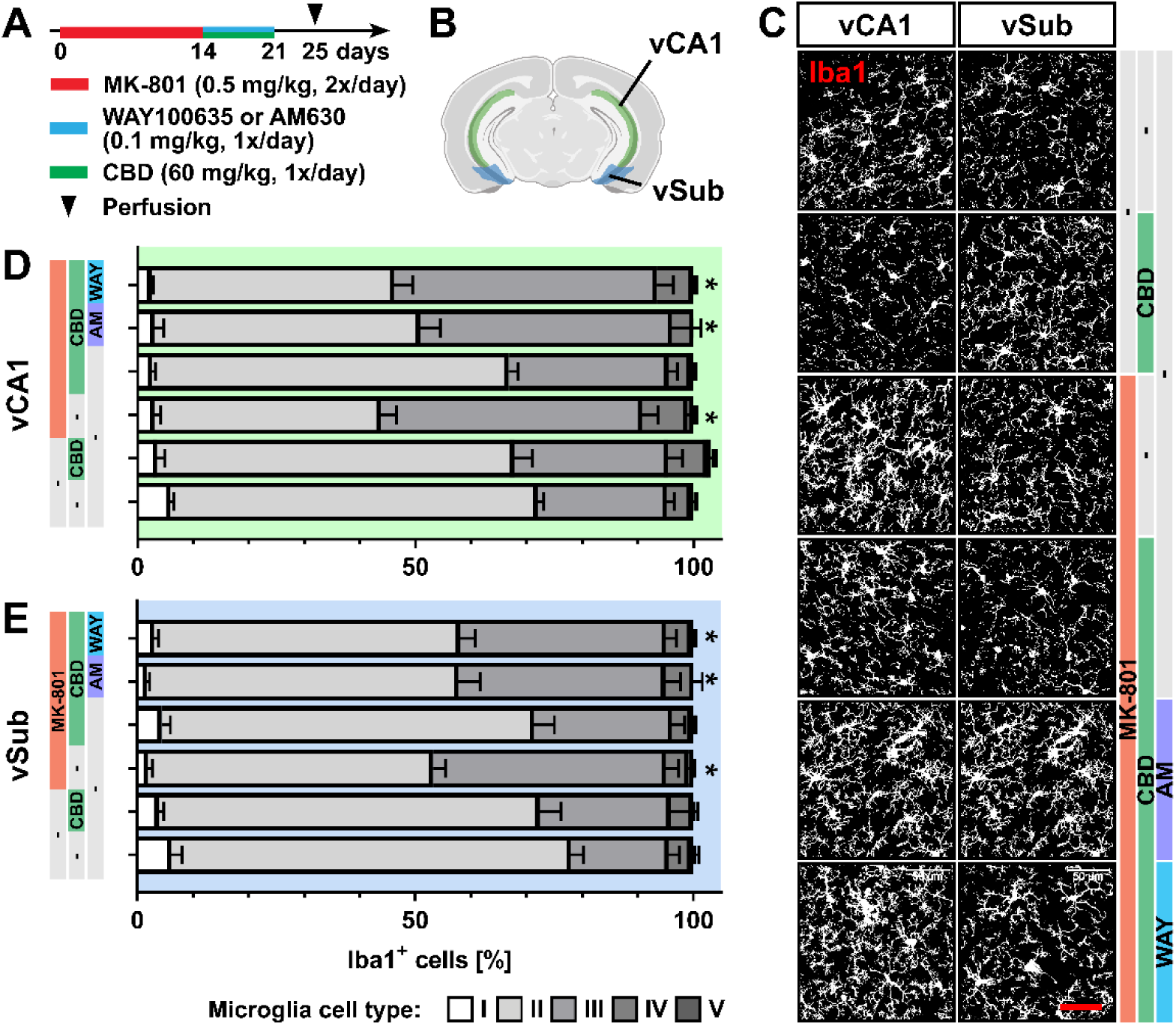
CBD reversed MK-801-induced microglia activation through 5-H1A and CB2 receptors in the hippocampus. **(A)** Experimental protocol for pharmacological treatment with MK-801, WAY100635, or AM630 and CBD followed by perfusion and immunohistochemical analysis. **(B)** Schematic representation of the ventral CA1 (vCA1) and the ventral subiculum (vSub) of the hippocampus used for analysis [Created with BioRender.com]. **(C)** Representative view of Iba1^+^ microglia. Scale bar, 50 µm. **(D-E)** Relative number of microglia divided into five classes (resting microglia, type I-II; reactive microglia, type III-V) in the vCA1 **(D)** and vSub **(E)** regions. Data are represented as mean ± SEM and derived from N=7-8 mice. Each group was compared to the control group (*, p < 0.05). Details on statistical analysis can be found in Supplementary Table 1.

MK-801 did not change the total density of microglial cells in any of the analyzed regions, suggesting that microglia proliferation and survival are unaffected (Supplementary Figure 5C-F). These findings indicate that CBD reverts MK-801-induced microglia activation in the mPFC and vHip by direct or indirect activation of both 5-HT1A and CB2 receptors.

## Discussion

The results of the present study corroborate previous works showing that CBD causes antipsychotic-like effects in animal models based on the chronic blockade of NMDAR [36, 37].

Repeated administration of the NMDAR antagonist MK-801 impaired novel object recognition, a test used to assess cognitive symptoms in SCZ models [37]. Although other studies have already shown that MK-801 impairs performance in the NOR test [9, 37, 49], long-term memory deficits observed one week after the end of treatment, as in our work, have yet to be fully explored in the literature. This cognitive impairment could depend on NMDAR antagonism interference on long-term potentiation (LTP) in the hippocampus-prefrontal cortex (PFC) pathway and the vSub [50-53]. Corroborating this possibility, LTP impairment has been found in SCZ patients [54]. In addition to this potential mechanism, our study showed that the memory deficit was associated with decreased γ oscillation. Brain γ oscillations are strongly associated with sensory and cognitive performance. They are generated from the activity of PV^+^ interneurons, which control the excitatory/inhibitory balance and the rhythmicity of the firing of pyramidal neurons in the prefrontal cortex [12, 14]. Our results are in line with previous research showing impairment in γ oscillation associated with SCZ and with SCZ-induced cognitive tasks [17, 55]. Although cortical γ oscillations are generated from the prefrontal cortex, we detected the reduction using whole-brain interhemispheric recordings acquired through depth electrodes implanted into the brain occipital lobes. More precise recording using microelectrodes implanted in the prefrontal cortex would allow the investigation of the mechanisms involved in generating γ oscillations in response to MK-801.

In line with previous studies [36, 56-58], we showed that MK-801 treatment decreased the number of PV^+^ cells in the mPFC and hippocampus. NMDAR antagonism affects GABAergic interneurons due to their high firing rate, which keeps them depolarized, making NMDAR on these neurons more likely to overcome Mg^+2^ blockade and thus be more vulnerable to these antagonists [59-61]. In this study, we observed a decrease in the number of PV^+^ cells in the prelimbic but not in the infralimbic regions of mPFC. These changes align with results observed after chronic treatment with phencyclidine, another NMDAR antagonist [62].

Several studies have shown that reduced PFC inhibitory transmission induces various cognitive, emotional, and dopaminergic abnormalities related to SCZ [63]. Reduced GABAergic activity in the PFC of rats decreases information processing speed, cognitive flexibility, retrieval of relevant information, and increases dopaminergic tone in the ventral tegmental area (VTA) [64, 65]. Few studies, however, have explored the differences in GABAergic changes in animal models of SCZ individualizing the prelimbic and infralimbic cortices. These regions, nonetheless, have opposing effects on the VTA dopamine system—prelimbic and infralimbic activation increases and decreases, respectively, VTA dopamine neuron activity [66]. In addition, opposite effects of interference on prelimbic and infralimbic cortices have been described on working memory [67-73].

In our study, MK-801 treatment also decreased PV^+^ cells in the vCA1 and vSub of the hippocampus. Several studies have found a decrease in these neurons in the hippocampus after treatment with NMDAR antagonists [18], although few have explored its ventral regions [51, 74].

Increased activity in the hippocampus is described in SCZ. This effect has been associated with functional loss of PV^+^ interneurons on pyramidal neurons [75-79]. By interfering with a circuit comprising the nucleus accumbens and ventral pallidum, this latter effect would increase the activity of VTA dopamine neurons, leading to the *hyperdopaminergia* in SCZ patients [80].

Changes in PNNs surrounding PV^+^ interneurons were observed in SCZ models using NMDAR antagonists [81-83]. We showed that MK-801 decreased PNNs around PV^+^ cells in the prelimbic mPFC and vHip and that CBD treatment reverted these changes. Our analysis benefits from the fact that the density of PNNs was evaluated around each PV^+^ cell surrounded by PNNs. In contrast, previous studies reported the number of PV^+^/PNN^+^ cells, which is inevitably affected by the reduction in the total density of PV^+^ cells.

In line with our results, alterations in PNNs co-localized with PV^+^ interneurons were observed in samples from SCZ patients [84-86]. Given the high metabolic demand of PV^+^ interneurons, deficits of PNNs could result in enhanced exposure to oxidative stress [87, 88] and decreased expression of PV^+^ cells [28]. Degradation of PNNs could alter the excitatory-inhibitory balance and γ oscillation [83, 89]. Therefore, the pro-cognitive effect of CBD in our study may be due to a restored excitatory-inhibitory balance resulting from the enhanced expression of PNNs on PV^+^ interneurons.

The memory impairment caused by MK-801 and the decreased number of PV^+^ cells and PNNs was associated with microglial activation. Several studies using NMDAR antagonists suggest an association between microglia activation and negative symptoms and cognitive deficits in SCZ [90-96]. Second-generation antipsychotics, including clozapine [97] and amisulpride [98], can reverse microglial activation. Also, co-treatment with the microglia inhibitor minocycline [90, 99], as well as non-steroidal anti-inflammatory drugs [100, 101], enhanced the effect of antipsychotic treatment of SCZ patients. Minocycline treatment attenuated the decrease in PV^+^ cells in neurodevelopmental disruption models of SCZ based on immune maternal activation [102]. In previous work, we demonstrated that CBD prevents, when administered concomitantly, MK-801-induced microglial activation [9]. In the present study, we expand this result by showing that CBD also reverses this effect since it was administered after the end of MK-801 treatment.

We also investigated the possible pharmacological mechanism responsible for the antipsychotic effects of CBD. CBD has a complex pharmacology, with more than 60 molecular targets [103]. Here, we focus on two potential targets, 5-HT1A and CB2 receptors, already associated with CBD antipsychotic action [37, 104].

CBD attenuated the effects of repeated MK-801 administration on short- and long-term memory via 5-HT1A receptors. The effects of CBD on short-term memory in the NOR test are already well described in several models of SCZ [9, 37, 105] and after THC administration on long-term memory [106]. By itself, CBD did not interfere with short- and long-term memory, corroborating findings in the water maze Morris test [107].

Although initially described as a 5-HT1A full agonist [108], CBD probably acts as a positive allosteric modulator of these receptors [109]. The involvement of serotonergic pathways in CBD-mediated effects corroborates results observed with atypical antipsychotics, such as aripiprazole, clozapine, lurasidone, tandospirone, and ziprasidone, which also act as partial agonists of 5-HT1A receptors [110-112].

5-HT1A receptors regulate neurogenesis, synaptogenesis, and the excitation/inhibition balance [113-116]. In the hippocampus, 5-HT levels are reduced in several preclinical SCZ models [117-120]. Therefore, CBD could compensate for these reduced 5-HT levels by facilitating the activation of 5-HT1A receptors expressed, among others, on PV^+^ interneurons [108].

In the GABAergic interneurons of the PFC and hippocampus, a high density of 5-HT1A receptors is responsible for neuronal hyperpolarization [121] and consequent disinhibition of pyramidal neurons in the hippocampus, which in turn regulate γ oscillations [122]. As already discussed, CBD attenuated the γ wave impairment induced by MK-801. These oscillations are closely associated with the activity of PV^+^ GABAergic interneurons [123]. Therefore, our findings suggest that CBD could re-establish GABAergic modulation on pyramidal neurons and restore the excitatory-inhibitory balance. This protective effect of CBD has been described before. It attenuated the depletion of PV^+^ interneurons in the hippocampus, but not in the PFC, in SCZ models based on neurodevelopmental disruption [48, 124]. Also, CBD decreased the atrophy and death of PV^+^ interneurons in the kainic acid model of temporal lobe epilepsy [125]. The protective effect of CBD on PV^+^ interneurons seem to depend on 5-HT1A and CB2 receptors in the vHIP but not in the prelimbic mPFC. At the moment, we have no explanation for this difference. A study with local CBD microinjection into the prelimbic cortex indicated that 5-HT1A receptors play a complex role in this structure [126]. CBD could produce anxiolytic or anxiogenic responses, depending on previous exposure to restraint stress and glucocorticoid levels. Both responses, however, were blocked by WAY100635. Further studies are needed to understand the role of these receptors and the mechanisms of the PV^+^/PNNs CBD protective effect in this region.

Our results also suggest that the endocannabinoid system could be involved in the protective effect of CBD on PV^+^ interneurons. Nevertheless, no evidence indicates that CB2 receptors are expressed in PV^+^ interneurons, suggesting the engagement of indirect mechanisms such as neuroinflammation inhibition. Microglial activation results in PNN degradation [127] and is observed in patients with SCZ [128-131]. In our study, CBD attenuation of the microglial activation induced by MK-801 was blocked by pretreatment with AM630 or WAY100635, indicating that this effect depends on 5-HT1A or CB2 receptors. Both receptors have been associated with inflammatory responses. 5-HT enhances microglia’s injury-induced mobility, phagocytic activity, and exosome secretion [132-134]. Resting microglia express CB1 and CB2 receptors at low levels. Upon activation, microglia significantly increase the synthesis of endocannabinoids and up-regulate the expression of CB2 receptors, whose activation increases the production of microglia-derived neuroprotective factors and reduces the production of pro-inflammatory factors [135, 136].

As previously discussed, CBD inhibits the FAAH enzyme, increasing levels of AEA [39]. In this perspective, considering that microglia produce approximately 20 times more endocannabinoids than neurons and other glial cells *in vitro* and may be the primary source of endocannabinoids in neuroinflammatory conditions [136, 137], the anti-inflammatory role of CBD may result from increased levels of available AEA and subsequent CB2 receptor activation.

In conclusion, this study described the significant effects of CBD on the behavioral and electrophysiological changes observed in animal models of SCZ. It also revealed that CBD targets critical processes associated with the development of this disorder, namely microglia activation and the decreased expression of PNNs and PV^+^ cells in the mPFC and vHip. Finally, we also showed that part of these effects depends on facilitating 5-HT1A and CB2 receptors’ activity.

## Acknowledgments

We thank Marcos Antonio de Carvalho and Eleni Taburus Gomes for their technical support. We thank Dr. Alline Cristina Campos and Dr. Mariza Bortolanza for helping with the confocal microscope analysis and the Wisteria Floribunda lectin assay, respectively. The authors thank the members of the Department of Molecular Physiology of the University of Saarland for their intellectual input, as well as Daniel Schauenburg and colleagues for their expert mouse maintenance.

## Supplementary Figures

**Supplementary Figure 1.**
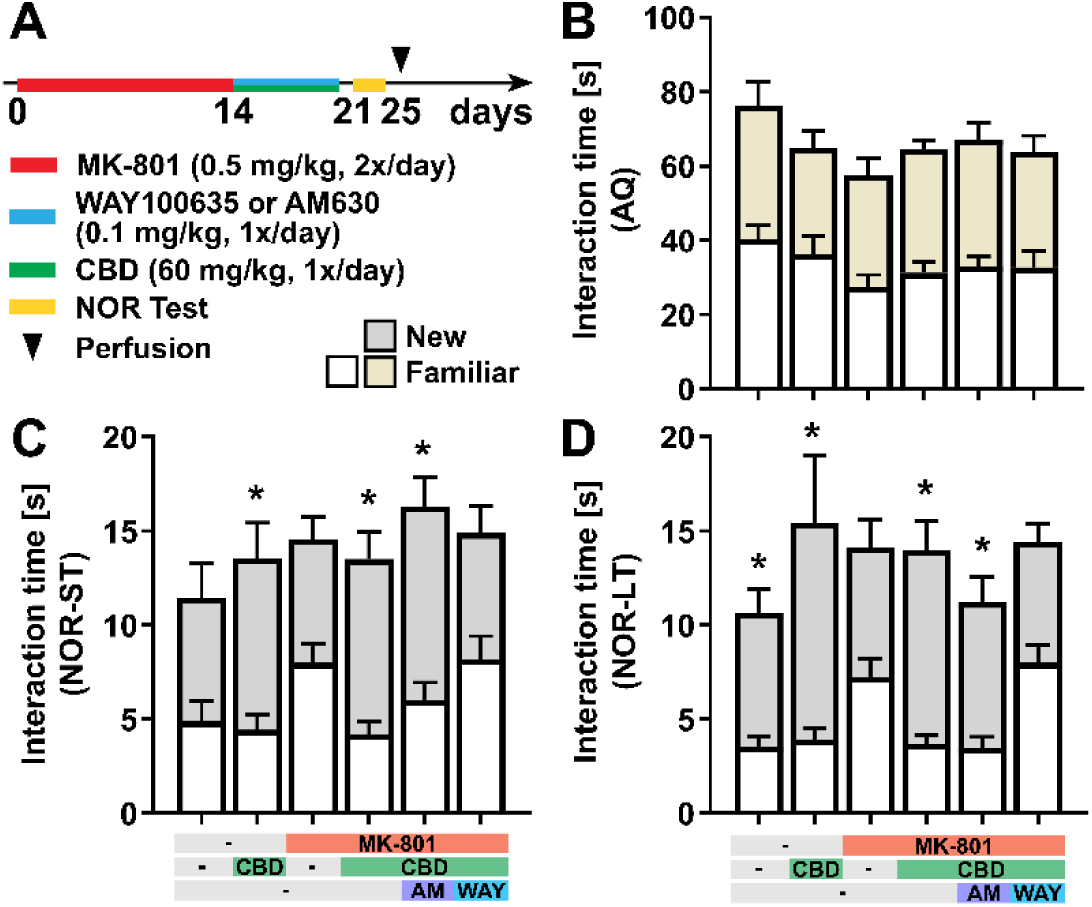
CBD treatment reduces MK-801-induced working memory impairment through 5-HT1A receptors. **(A)** Experimental protocol for pharmacological treatment with the NMDA receptor antagonist MK-801, WAY100635, or AM630 and CBD followed by novel-object-recognition test. **(B-D)** Absolute exploration time with the new and familiar objects during the acquisition phase (AQ, B), the short-term memory retention phase (NOR-ST, C), and the long-term memory retention phase (NOR-LT, **D**). Data are represented as mean ± SEM and derived from N=8-9 mice. Details on statistical analysis can be found in Supplementary Table 1. For the group, the interaction time with the two objects was compared (*, p < 0.05).

**Supplementary Figure 2.**
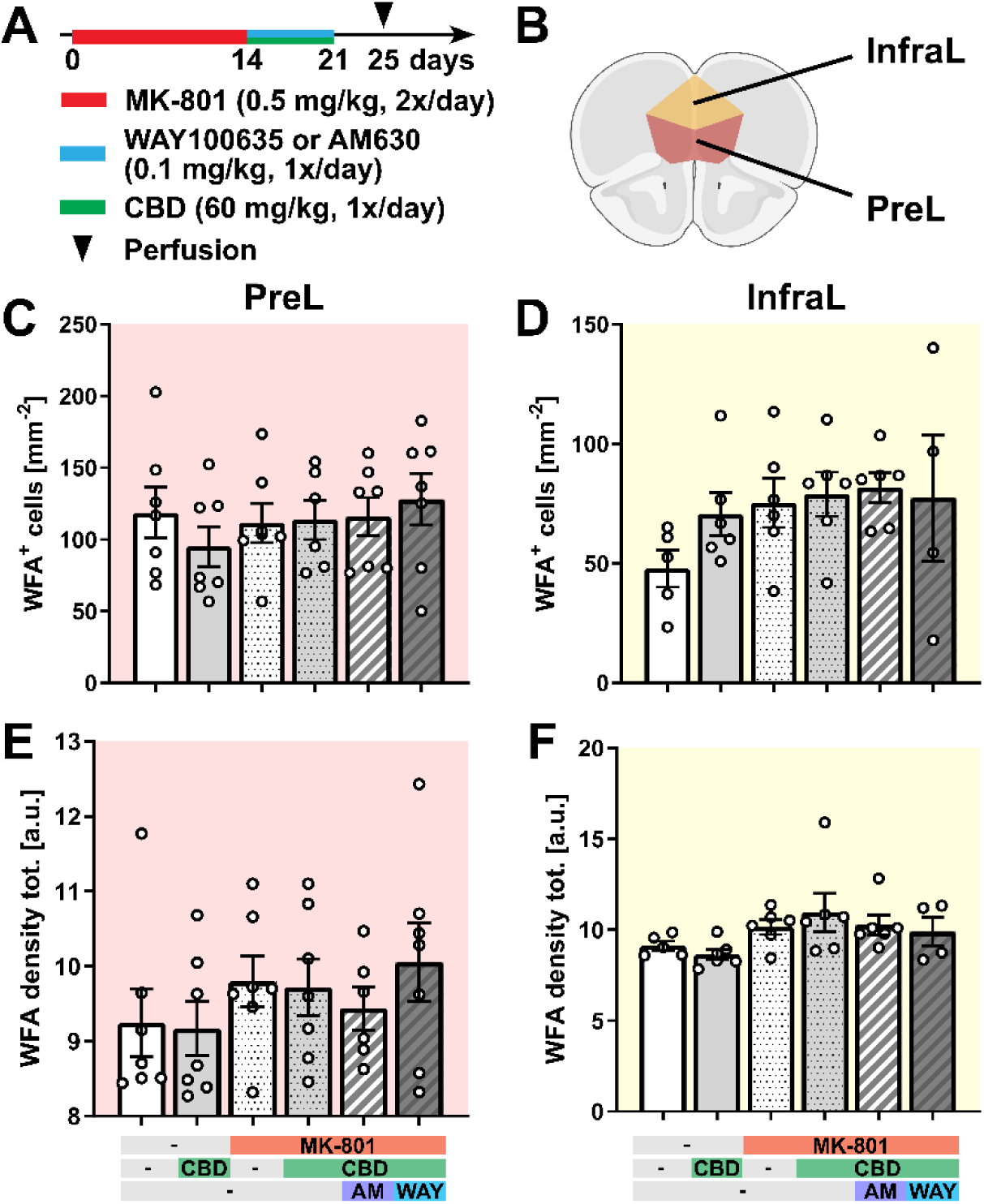
There is no difference in WFA^+^ cell density and WFA density across the pharmacological treatments in the mPFC. **(A)** Experimental protocol for pharmacological treatment with MK-801, WAY100635, or AM630 and CBD followed by perfusion and immunohistochemical analysis. **(B)** Schematic representation of the prelimbic region (PreL) and the infralimbic region (InfraL) of the medial prefrontal cortex (mPFC) used for analysis [Created with BioRender.com]. **(C-D)** Total cell density of WFA^+^ cells in the PreL **(C)** and InfraL **(D)** regions. **(E-F)** Total WFA density (calculated as fluorescence intensity) in the PreL **(E)** and InfraL **(F)** regions. Data are represented as mean ± SEM and derived from N=4-7 mice. Details on statistical analysis can be found in Supplementary Table 1.

**Supplementary Figure 3.**
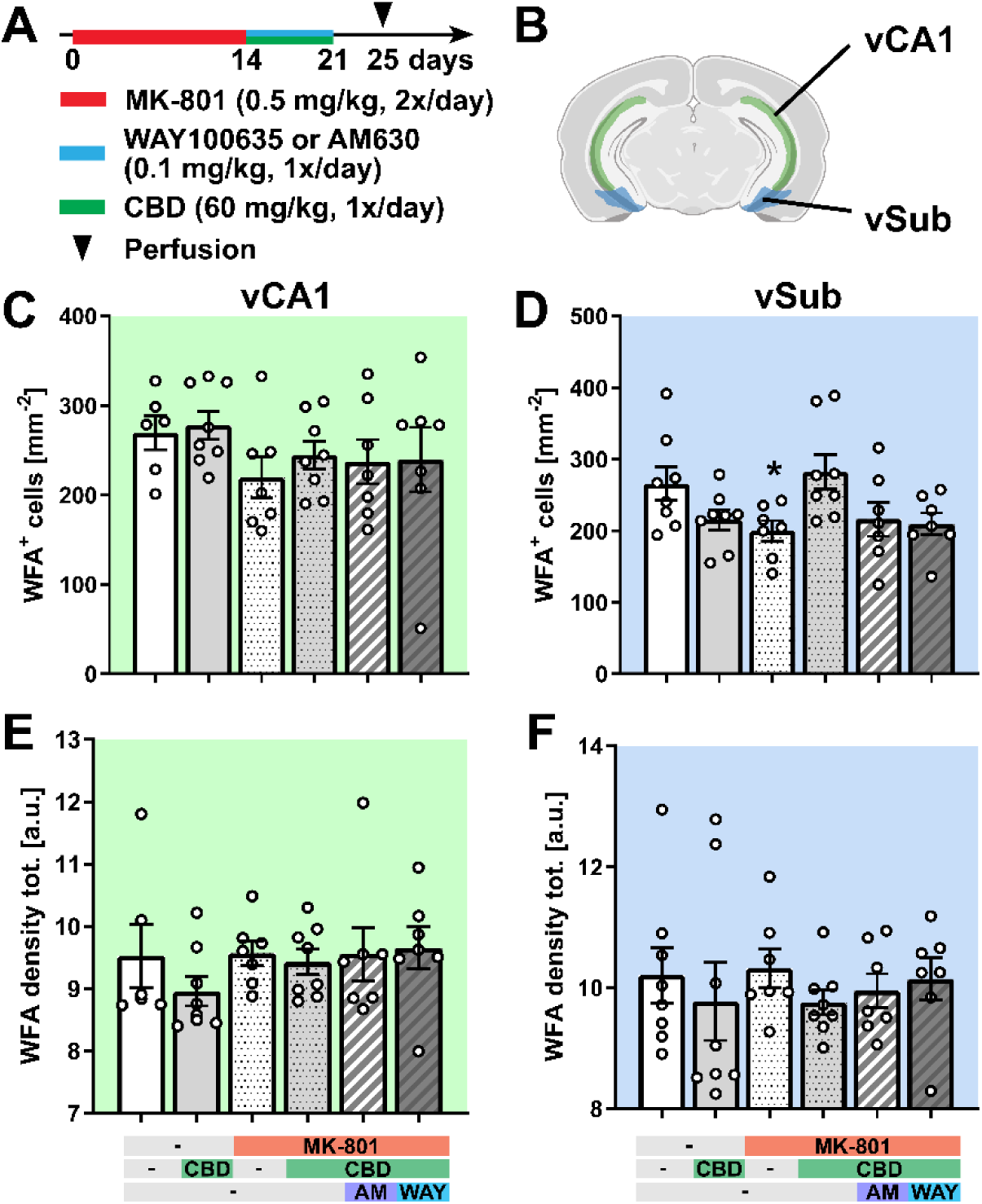
There is no difference in WFA^+^ cell density and WFA density across the pharmacological treatments in the hippocampus. **(A)** Experimental protocol for pharmacological treatment with MK-801, WAY100635, or AM630 and CBD followed by perfusion and immunohistochemical analysis. **(B)** Schematic representation of the ventral CA1 (vCA1) and the ventral subiculum (vSub) of the hippocampus used for analysis [Created with BioRender.com]. **(C-D)** Total cell density of WFA^+^ cells in the vCA1 **(C)** and vSub **(D)** regions. **(E-F)** Total WFA density (calculated as fluorescence intensity) in the vCA1 **(E)** and vSub **(F)** regions. Data are represented as mean ± SEM and derived from N=6-9 mice. Details on statistical analysis can be found in Supplementary Table 1.

**Supplementary Figure 4.**
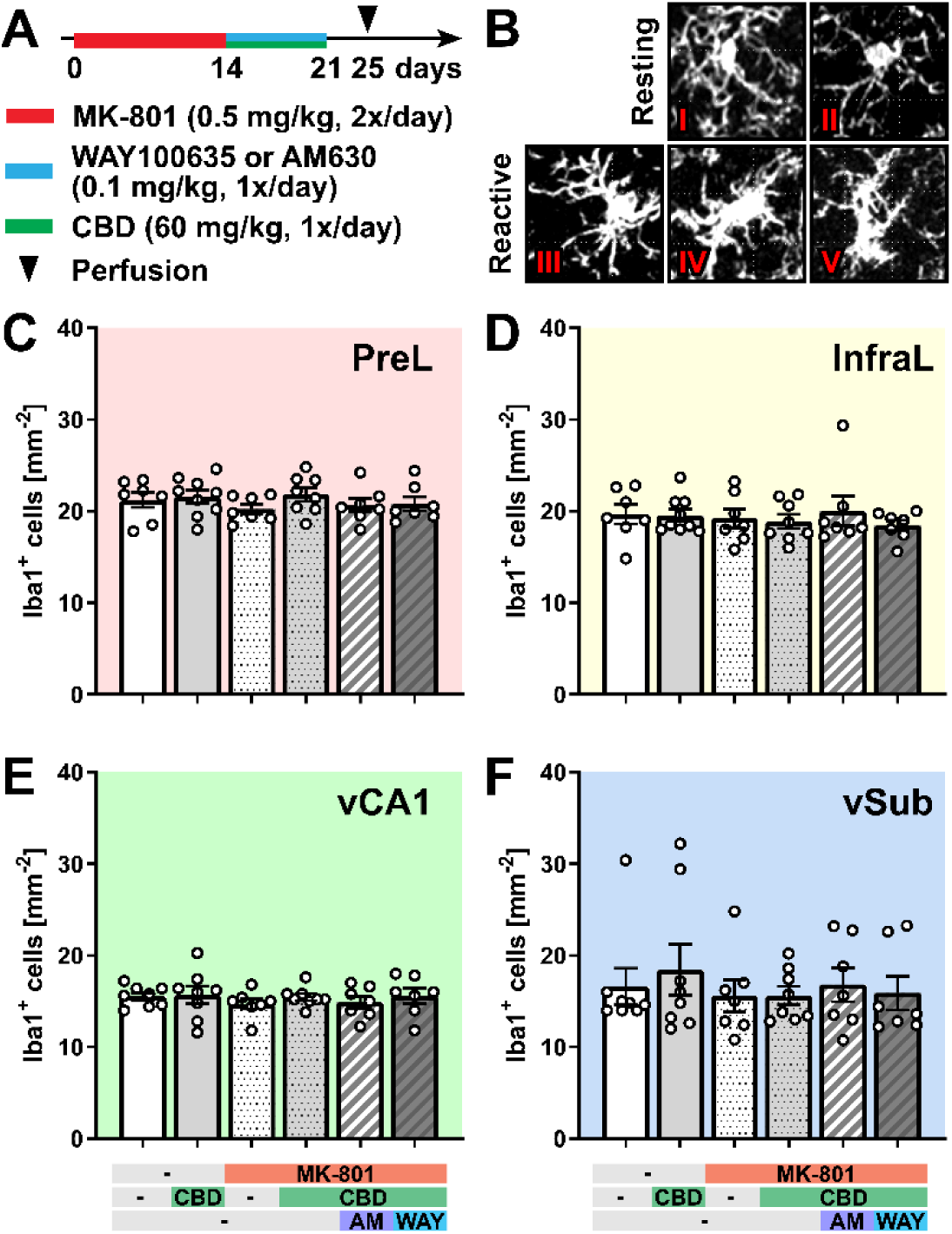
There is no difference in total Iba1^+^ cell density across the pharmacological treatments. **(A)** Experimental protocol for pharmacological treatment with MK-801, WAY100635, or AM630 and CBD followed by perfusion and immunohistochemical analysis. **(B)** Schematic representation of the morphological classification of microglial cells into five classes: resting microglia (type I-II) and reactive microglia (type III-V). **(C-F)** Total cell density of Iba1^+^ cells in the PreL **(C)** and InfraL region **(D)** of the mPFC, as well as the vCA1 **(E)** and vSub **(F)** regions of the hippocampus. Data are represented as mean ± SEM and derived from N=7-9 mice. Details on statistical analysis can be found in Supplementary Table 1.

**Supplementary Table 1.**

| Figure | Levene's homogeneity test | Statistical test | F | p-value | Multiple comparisons |  |
| --- | --- | --- | --- | --- | --- | --- |
| **Fig.1B** | $p > 0.05$ | one-way ANOVA with Dunnett post-hoc test | 9.833 | $p < 0.0001$ | Amph<br>Amph-CBD<br>Amph-CBD-AM630<br>Amph-CBD-WAY | $p = 0.0001$<br>$p = 0.1485$<br>$p = 0.0001$<br>$p = 0.0034$ |
| **Fig.1D** | $p < 0.05$ | one-way ANOVA | 0.937 | $p = 0.465$ | | |
| **Fig.1E** | $p > 0.05$ | one-way ANOVA with Dunnett post-hoc test | 8.348 | $p < 0.0001$ | CBD<br>MK-801<br>MK-801-CBD<br>MK-801-CBD-AM630<br>MK-801-CBD-WAY | $p = 0.9911$<br>$p = 0.0078$<br>$p = 0.8220$<br>$p = 0.9999$<br>$p = 0.0024$ |
| **Fig.1F** | $p < 0.05$ | one-way ANOVA with Dunnett post-hoc test | 7.177 | $p < 0.0001$ | CBD<br>MK-801<br>MK-801-CBD<br>MK-801-CBD-AM630<br>MK-801-CBD-WAY | $p = 0.9986$<br>$p = 0.0237$<br>$p = 0.8061$<br>$p = 0.9996$<br>$p = 0.0049$ |
| **Fig.3D** | $p \leq 0.05$ | one-way ANOVA with Dunnett post-hoc test | 5.126 | $p = 0.0012$ | CBD<br>MK-801<br>MK-801-CBD<br>MK-801-CBD-AM630<br>MK-801-CBD-WAY | $p = 0.0146$<br>$p = 0.0002$<br>$p = 0.3367$<br>$p = 0.2713$<br>$p = 0.6203$ |
| **Fig.2C** | - | one-way ANOVA with Dunnett post-hoc test | 12.01 | $p = 0.0039$ | MK-801<br>MK-801-CBD | $p = 0.0119$<br>$p = 0.4173$ |
| **Fig.3E** | $p \leq 0.05$ | one-way ANOVA | 0.2272 | $p = 0.9483$ | | |
| **Fig.3F** | $p \leq 0.05$ | one-way ANOVA with Dunnett post-hoc test | 2.378 | $p = 0.0580$ | CBD<br>MK-801<br>MK-801-CBD<br>MK-801-CBD-AM630<br>MK-801-CBD-WAY | $p = 0.0647$<br>$p = 0.0500$<br>$p = 0.8062$<br>$p = 0.5494$<br>$p = 0.9955$ |
| **Fig.3G** | $p > 0.05$ | one-way ANOVA | 0.6096 | $p = 0.6930$ | | |
| **Fig.3H** | $p > 0.05$ | one-way ANOVA with S-N-K post-hoc test | 4.555 | $p = 0.0004$ | CBD<br>MK-801<br>MK-801-CBD<br>MK-801-CBD-AM630<br>MK-801-CBD-WAY | MD = -0.0031<br>MD = 0.01299<br>MD = -0.0124<br>MD = -0.0057<br>MD = -0.0070 |
| **Fig.3I** | $p > 0.05$ | one-way ANOVA | 0.4595 | $p = 0.8064$ | | |
| **Fig.4D** | $p \leq 0.05$ | one-way ANOVA with Dunnett post-hoc test | 6.060 | $p = 0.0003$ | CBD<br>MK-801<br>MK-801-CBD<br>MK-801-CBD-AM630<br>MK-801-CBD-WAY | $p = 0.2795$<br>$p = 0.0002$<br>$p = 0.0936$<br>$p = 0.0014$<br>$p = 0.0059$ |
| **Fig.4E** | $p > 0.05$ | one-way ANOVA with Dunnett post-hoc test | 5.790 | $p = 0.0004$ | CBD<br>MK-801<br>MK-801-CBD<br>MK-801-CBD-AM630<br>MK-801-CBD-WAY | $p = 0.7640$<br>$p = 0.0021$<br>$p = 0.7722$<br>$p = 0.0020$<br>$p = 0.0085$ |
| **Fig.4F** | $p > 0.05$ | one-way ANOVA with Dunnett post-hoc test | 4.716 | $p = 0.0020$ | CBD<br>MK-801<br>MK-801-CBD<br>MK-801-CBD-AM630<br>MK-801-CBD-WAY | $p = 0.9676$<br>$p = 0.0047$<br>$p = 0.8534$<br>$p = 0.0477$<br>$p = 0.0485$ |
| **Fig.4G** | $p \leq 0.05$ | one-way ANOVA with Dunnett post-hoc test | 6.476 | $p = 0.0002$ | CBD<br>MK-801<br>MK-801-CBD<br>MK-801-CBD-AM630<br>MK-801-CBD-WAY | $p = 0.4490$<br>$p = 0.0020$<br>$p = 0.7551$<br>$p = 0.0021$<br>$p = 0.0007$ |
| **Fig.4H** | $p > 0.05$ | one-way ANOVA with Dunnett post-hoc test | 6.059 | $p < 0.0001$ | CBD<br>MK-801<br>MK-801-CBD<br>MK-801-CBD-AM630<br>MK-801-CBD-WAY | $p = 0.3390$<br>$p < 0.0001$<br>$p = 0.3959$<br>$p = 0.1419$<br>$p = 0.0099$ |
| **Fig.4I** | $p \leq 0.05$ | one-way ANOVA<br>with Dunnett post-hoc test | 16.62 | $p < 0.0001$ | CBD<br>MK-801<br>MK-801-CBD<br>MK-801-CBD-AM630<br>MK-801-CBD-WAY | $p = 0.6991$<br>$p < 0.0001$<br>$p = 0.3599$<br>$p = 0.0148$<br>$p < 0.0001$ |
| **Fig.5D** | $p < 0.05$ | one-way ANOVA<br>with Dunnett post-hoc test | 7.254 | $p < 0.0001$ | CBD<br>MK-801<br>MK-801-CBD<br>MK-801-CBD-AM630<br>MK-801-CBD-WAY | $p = 0.9813$<br>$p = 0.0045$<br>$p = 0.9201$<br>$p = 0.0497$<br>$p = 0.0774$ |
| **Fig.5E** | $p > 0.05$ | one-way ANOVA<br>with Dunnett post-hoc test | 12.508 | $p < 0.0001$ | CBD<br>MK-801<br>MK-801-CBD<br>MK-801-CBD-AM630<br>MK-801-CBD-WAY | $p = 0.9811$<br>$p = 0.0020$<br>$p = 0.8831$<br>$p = 0.0015$<br>$p = 0.0010$ |
| **Fig.5D** | $p > 0.05$ | one-way ANOVA<br>with Dunnett post-hoc test | 16.603 | $p < 0.0001$ | CBD<br>MK-801<br>MK-801-CBD<br>MK-801-CBD-AM630<br>MK-801-CBD-WAY | $p = 0.2520$<br>$p < 0.0001$<br>$p = 0.5762$<br>$p < 0.0001$<br>$p < 0.0001$ |
| **Fig.5E** | $p > 0.05$ | one-way ANOVA<br>with Dunnett post-hoc test | 10.340 | $p < 0.0001$ | CBD<br>MK-801<br>MK-801-CBD<br>MK-801-CBD-AM630<br>MK-801-CBD-WAY | $p = 0.5709$<br>$p < 0.0001$<br>$p = 0.4258$<br>$p = 0.0002$<br>$p = 0.0003$ |
| **Suppl.**<br>**Fig.1B** | - | Unpaired t-test | - | - | VEH<br>CBD<br>MK-801<br>MK-801-CBD<br>MK-801-CBD-AM630<br>MK-801-CBD-WAY | $p = 0.5638$<br>$p = 0.2230$<br>$p = 0.0894$<br>$p = 0.0844$<br>$p = 0.1622$<br>$p = 0.5219$ |
| **Suppl.**<br>**Fig.1C** | - | Unpaired t-test | - | - | VEH<br>CBD<br>MK-801<br>MK-801-CBD<br>MK-801-CBD-AM630<br>MK-801-CBD-WAY | $p = 0.4497$<br>$p = 0.0371$<br>$p = 0.6444$<br>$p = 0.0046$<br>$p = 0.0301$<br>$p = 0.4365$ |
| **Suppl.**<br>**Fig.1D** | - | Unpaired t-test | - | - | VEH<br>CBD<br>MK-801<br>MK-801-CBD<br>MK-801-CBD-AM630<br>MK-801-CBD-WAY | $p = 0.0195$<br>$p = 0.0508$<br>$p = 0.6935$<br>$p = 0.0006$<br>$p = 0.0098$<br>$p = 0.2546$ |
| **Suppl.**<br>**Fig.2C** | $p > 0.05$ | one-way ANOVA | 0.5146 | $p = 0.7634$ | - | - |
| **Suppl.**<br>**Fig.2D** | $p > 0.05$ | one-way ANOVA | 1.126 | $p = 0.3705$ | - | - |
| **Suppl.**<br>**Fig.2E** | $p > 0.05$ | one-way ANOVA | 0.8872 | $p = 0.4995$ | - | - |
| **Suppl.**<br>**Fig.2F** | $p > 0.05$ | one-way ANOVA | 1.403 | $p = 0.2466$ | - | - |
| **Suppl.**<br>**Fig.3C** | $p > 0.05$ | one-way ANOVA | 0.7402 | $p = 0.5985$ | - | - |
| **Suppl.**<br>**Fig.3D** | $p > 0.05$ | one-way ANOVA | 1.862 | $p = 0.1343$ | - | - |
| **Suppl.**<br>**Fig.3E** | $p > 0.05$ | one-way ANOVA | 0.6669 | $p = 0.6509$ | - | - |
| **Suppl.**<br>**Fig.3F** | $p \leq 0.05$ | one-way ANOVA | 0.3290 | $p = 0.8924$ | - | - |
| **Suppl.**<br>**Fig.4C** | $p > 0.05$ | one-way ANOVA | 0.6895 | $p = 0.6343$ | - | - |
| **Suppl.**<br>**Fig.4D** | $p > 0.05$ | one-way ANOVA | 0.3293 | $p = 0.8922$ | - | - |
| **Suppl.**<br>**Fig.4E** | $p > 0.05$ | one-way ANOVA | 0.3696 | $p = 0.8663$ | - | - |
| **Suppl.**<br>**Fig.4F** | $p > 0.05$ | one-way ANOVA | 0.3134 | $p = 0.9019$ | - | - |

